# Age-dependent phosphorylation of estrogen receptor beta (ERβ) regulates homodimer formation and impairs the repression of amyloid precursor protein (APP) promoter

**DOI:** 10.64898/2026.11.13.766544

**Authors:** Yoldas Yildiz, Jaroslava Seflova, Sarah Flury, Destiny Deschenes, Seth L. Robia, Toni R. Pak

**Affiliations:** Department of Cell & Molecular Physiology, Stritch School of Medicine, Loyola University Chicago, Maywood, IL, USA

## Abstract

Estrogen receptor beta (ERβ) is a ligand-activated nuclear steroid receptor that promotes neuroprotection through its transcriptional coordination of "healthy cell" gene programs. However, circulating estrogens are dramatically reduced post-menopause and the administration of hormone replacement therapy is only beneficial within a narrowly restricted timeframe after the menopause transition. The molecular underpinnings for this failure of ERβ to evoke protection in advanced age is unknown. Evidence suggests that age-related changes in protein–protein interactions could preclude ERβ genomic signaling. Protein–protein interactions can be regulated and modified through post-translational modifications (PTM)s of individual amino acid residues; the most well-studied PTM being phosphorylation. Our prior work showed that ERβ phosphorylation was age-dependent leading to the hypothesis that ERβ phosphorylation is required for ERβ–protein interactions, specifically ERβ homodimer formation. We used fluorescence lifetime measurements to quantify Förster resonance energy transfer (FRET) of ERβ-S176, S200, and Y488 phosphomutant protein interactions. Overall, our results demonstrate that phosphorylation of ERβ-S176, S200, and Y488 reduced ERβ homodimer formation, suggesting an age-dependent molecular mechanism that regulates ERβ genomic signaling. Moreover, phosphorylation at S200 and Y488 resulted in a loss of ERβ-mediated amyloid precursor protein (APP) promoter gene repression providing evidence that these age-dependent PTMs directly impact ERβ’s function as a transcription factor. Taken together, these data suggest that age-dependent changes in ERβ phosphorylation result in decreased ERβ genomic signaling, which could reduce the neuroprotective effects of estrogens following menopause.

**SIGNIFICANCE STATEMENT:** Our findings demonstrate that age-dependent post-translational modifications of ERβ could mechanistically explain the timing paradox observed with hormone replacement therapy in post-menopausal women by altering ERβ–protein interactions that are critical for mediating ERβ signaling pathways. Phosphorylation in the ERβ DNA binding domain regulated ERβ homodimer formation, which was required for its transcriptional activity. This domain is highly conserved among all members of the nuclear receptor family making these findings broadly applicable.

## INTRODUCTION

A major challenge in recent years has been understanding why hormone replacement therapy is less effective in late menopausal women [1-3]. Not only are the protective effects of estrogens diminished the longer women wait to initiate hormone therapy following menopause, but hormone administration in advanced age can increase the risk of major disease. Circulating estrogens drastically decline at menopause, which coincides with a risk of decreased cognitive function leading to the premise that estrogen treatment will restore optimal cognition. These neuroprotective effects of estrogens are mediated by estrogen receptors (ER)s, in part, by coordinating transcriptional gene programs that reduce oxidative stress, neuroinflammation, and maintain synaptic plasticity [4-6]. The two main estrogen receptors, ERα/ESR1 and ERβ/ESR2, are classified as ligand-activated transcription factors. They have many overlapping, but also distinct, functions due to differences in tissue distribution and abundance, structural elements, coregulatory protein recruitment and target gene selectivity [4, 7-9]. Post-translational modifications (PTM)s, such as phosphorylation, are an additional source of variation that contribute to the distinct functions of ERα and ERβ [10, 11]. Here, we functionally validated novel age–dependent ERβ phosphorylation sites that could mechanistically explain age-related shifts in the physiological response to estrogens.

Phosphorylation sites on ERβ have been experimentally validated primarily in the N-terminal intrinsically disordered AF–1 region. These sites are conserved between human, rat, and mouse and phosphorylation at several amino acids in the AF–1 domain confers ligand–independent activation of ERβ signaling including homodimer (ERβ–ERβ) formation, DNA binding, and recruitment of transcriptional co-activator proteins [12, 13]. However, there are predicted phosphorylation sites in other domains within ERβ that have yet to be experimentally tested, and none have been tested in an aging context, so their impact on age-dependent ERβ signaling is less clear. We previously identified two novel age-dependent phosphorylation sites (S176, S200) in the DNA binding domain of ERβ. Phosphomimetic manipulation at each of these sites resulted in decreased DNA binding and activation of gene transcription mediated by estrogen response elements (EREs) [14]. Moreover, using absolute quantification by mass spectrometry (AQUA–MS) we showed that phosphorylation of these sites decreased with age in the brains of women and in a rat model of menopause [14]. Our observation of age-related decreased phosphorylation at these sites suggests that there might be a post-menopausal shift in ERβ signaling away from gene programs that confer neuroprotection.

Classic genomic ERβ signaling is dependent on the ligand binding-induced formation of ERβ–ERβ homodimers, followed by subsequent engagement with chromatin and recruitment of transcriptional coregulators. Therefore, our goal in this study was to determine how phosphorylation at these previously uncharacterized sites affects ERβ receptor dimerization. We used Förster resonance energy transfer (FRET) combined with fluorescent lifetime imaging microscopy (FLIM) and showed that phosphorylation of S176, S200, or Y488 significantly decreased ERβ–ERβ dimerization. The effect of phosphorylation on these protein interactions was different in HEK293T cells compared to neuronal cells, underscoring a novel potential mechanism for the cell–type specificity of ERβ signaling. Overall, these studies add to our understanding of age–related molecular changes that directly alter ERβ function that could contribute to decreased estrogen-mediated neuroprotection in late menopause.

## RESULTS

### *ER*β–*ER*β affinity and homodimer formation was greater in neuronal cells (Fig. 1)

We measured ERβ–ERβ interactions using Förster resonance energy transfer (FRET) in two different human cell types: HEK293T and SK–N–SH neuronal cells. Briefly, we created expression vectors with unique fluorophores (mCyRFP1, mMaroon) at the N– or C–terminus of full–length human ERβ (530 aa, ERβ–WT). Quantification of FRET from multiexponential fitting of the fluorescence decay revealed several key parameters. The average fluorescence decay time (τ_avg_) was used to calculate average FRET efficiency where a lower τ_avg_ (i.e. faster donor decay rate) indicated higher average FRET efficiency. The calculated average FRET efficiency included the fluorophore decay rate of ERβ-donors that did not engage in FRET (i.e. ERβ monomers) as well as those that did engage in FRET (i.e. ERβ–ERβ dimers) (Fig. 1A). Using lifetime measurements of fluorescence, we then resolved subcomponents of τ_avg_ (τ_1_, τ_2_) to further explain changes in FRET. τ_1_ was the same fluorescence lifetime as the decay observed for cells expressing donor alone, so this parameter was taken to represent monomeric ERβ (Fig. 1A). τ_2_ was a shorter lifetime, taken to represent the intrinsic FRET of the ERβ-ERβ dimer complex containing a donor and acceptor. Thus, we used τ_2_ to calculate intrinsic FRET efficacy and the apparent donor-acceptor separation distance to gain insight into the overall conformation of the dimer (see Methods). The relative amplitudes of τ_2_ were used to determine the population of the bound FRET complexes (i.e. % dimer), with changes in τ_2_ amplitude taken to reflect changes in dimerization affinity (Fig. 1A).

**Figure 1.**
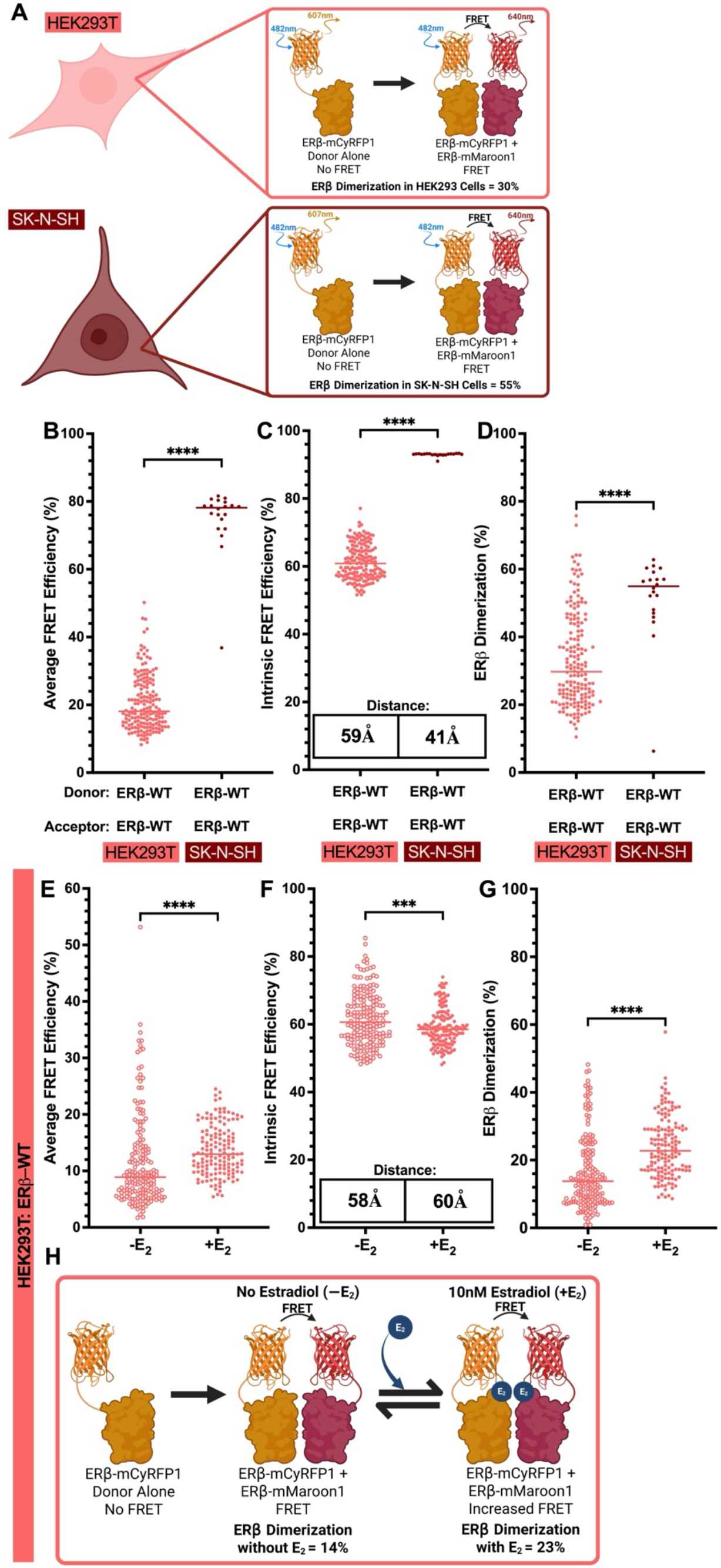
ERβ:ERβ affinity and homodimer formation was greater in neuronal cells. (A) Cartoon depiction of cell models, fluorescent tagged ERβ constructs with donor alone (no FRET) or dimerization (FRET). (B) Average FRET efficiency of all ERβ protomers, (C) intrinsic FRET efficiency of ERβ–ERβ dimers and calculated fluorophore distance (Å), and (D) percentage (%) dimer–formation was calculated using time-correlated single-photon counting (TCSPC) in HEK293T and SK–N–SH cells. (E) Average FRET efficiency of all ERβ protomers, (F) intrinsic FRET efficiency of ERβ–ERβ dimers and calculated fluorophore distance (Å), and (G) percentage (%) dimer–formation in HEK293T cells ± 10 nM E2 for 16 hours. (H) Cartoon depiction of quantified data from panels E–G. Data are displayed as median ± SD. Data were collected across 4 different biological replicates and 2 independent transfections per biological replicate. A non-parametric Kolmogorov Smirnov test was used for statistical analysis. An *** denotes p<0.001, **** p<0.0001. Created in BioRender. Yildiz, Y. (2026) https://BioRender.com/vfu6fcv and https://BioRender.com/yq6of1o

Consistent with known ERβ behavior, ERβ–WT formed homodimers and exhibited FRET in both cell types (Fig. 1B–D). However, our data demonstrated that the average FRET efficiency was significantly greater in neuronal cells compared to HEK293T cells (Fig. 1B). To determine whether this cell-type specific increase in average FRET efficiency was due to a structural change, or change in affinity, or both, we measured the amplitude of τ_2_ to determine the intrinsic FRET of the dimer complex and calculated the apparent fluorophore distance. Fluorophore distance was much farther apart in HEK293T cells (59 Å) compared to SK–N–SH cells (41 Å), consistent with a higher intrinsic FRET efficiency in the neuronal cells (Fig. 1C). Moreover, ERβ homodimer formation was greater in SK–N–SH neuronal cells (55%) compared to HEK293T cells (30%) (Fig. 1D) raising the possibility that neuronal cells express unique coregulatory proteins that facilitate ERβ–protein interactions.

It is well established that ERβ binding to ligands, such as 17β–estradiol (E2), facilitates dimer formation and induces a structural change in helix 12 (H12) that allows for coactivator protein interactions at the AF–2 groove [15-17]. Therefore, we next incubated HEK293T cells in media containing 10% charcoal-stripped FBS to remove all potential ERβ ligands, followed by treatment with ± 10 nM E2. Importantly, our initial FRET measurements in HEK293T cells were done in cells kept in regular growth media, which contained 10% FBS and therefore, sufficient levels of E2 and other growth factors to activate ERβ. Not surprisingly, average FRET efficiency was much lower in HEK293T cells grown in media containing charcoal-stripped FBS without E2 (Fig. 1E, -E2) and E2 supplementation restored the average FRET efficiency (Fig. 1E, +E2) comparable to that observed in regular growth media. On the other hand, there was a modest, yet significant, decrease in intrinsic FRET due to an increase in fluorophore distance (Fig. 1F). This was consistent with the expected E2–induced conformational change in ERβ. We also observed significantly fewer dimers in cells cultured in media with charcoal–stripped FBS without E2 compared to cells in regular media containing 10% FBS. For example, ERβ homodimer formation was 29.7% in regular HEK293T growth media (Fig. 1D), but only 13.7% in charcoal–stripped media, which then increased to 22.7% after 10 nM E2 treatment (Fig. 1G). Notably, ERβ dimer formation occurred in all media conditions, demonstrating that ERβ can readily form dimers even in the apo state.

### ERβ phosphorylation in the DNA binding domain decreased ERβ–ERβ interactions (*Fig. 2*)

We previously identified two age–dependent phosphosites (S176, S200) in the ERβ DNA binding domain (DBD) using a quantitative mass spectrometry approach (AQUA–MS) [14]. That study demonstrated that phosphorylation at ERβ–S176 and S200 significantly decreased with age in women’s brains and in the brain of a rat model of menopause. Prior to that study, these sites in the ERβ DBD had not been shown to be phosphorylated in any tissue. Therefore, our goal here was to extend those findings to better understand the functional consequences of age-dependent phosphorylation in the DBD. To test how phosphorylation in the DBD impacts the ability of ERβ to form homodimers, we used site-directed mutagenesis to create phosphonull (serine-to-alanine, S>A) and phosphomimetic (serine-to-glutamic acid, S>E) fluorescently tagged ERβ (Fig. 2A).

**Figure 2.**
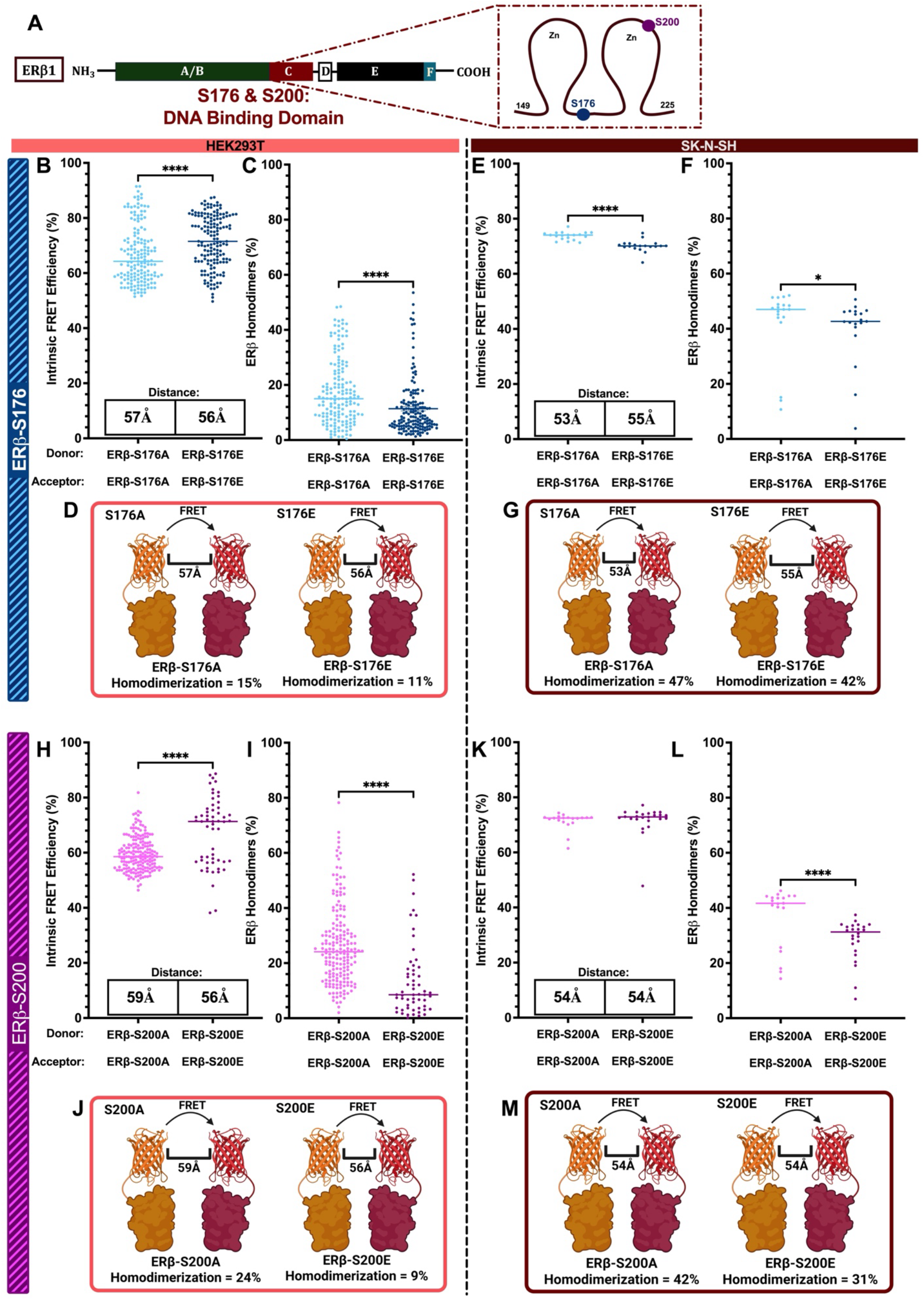
ERβ phosphorylation in the DNA binding domain decreased ERβ:ERβ interactions. (A) Diagram depicting ERβ protein domains. Expanded box of C-domain shows serine residues in the DNA binding domain that were mutated to phosponull (S>A) or phosphomimetic (S>E). Intrinsic FRET efficiency, fluorophore distance, and homodimer formation comparing S176A to S176E (blue) were calculated using time-correlated single-photon counting (TCSPC) in HEK293T (B, C) and SK-N-SH cells (E, F). Panels H, I and K, L show these same parameters for S200A/E (purple). Cartoon depictions of quantified data are illustrated in panels D, G for S176 and J, M for S200. Data are displayed as median ± SD. Data were collected across 4 different biological replicates and 2 independent transfections per biological replicate. A non-parametric Kolmogorov Smirnov test was used for statistical analysis. An * denotes p<0.05, **** p<0.0001. Created in BioRender. Yildiz, Y. (2026) https://BioRender.com/3hjibmy

Our results showed that intrinsic FRET efficiency was higher for ERβ–S176E due to a small structural shift that decreased fluorophore distance compared to ERβ–S176A in HEK293T cells (Fig. 2B). We also observed that, the phosphomimetic mutation resulted in significantly fewer dimers (Fig. 2C, D). By contrast, intrinsic FRET efficiency of ERβ–S176E decreased corresponding to a longer distance between fluorophores (Fig. 2E), and ERβ–S176E formed fewer dimers in SK–N–SH cells (Fig. 2F, G). These results suggested that phosphorylation of ERβ–S176 would induce cell–type specific structural changes and reduce ERβ–ERβ affinity.

ERβ–S200 is in the second zinc finger of the DBD and within a predicted "D box", which is a critical region for nuclear steroid receptor dimerization once bound to DNA [12]. Similar to what we observed for S176E, ERβ–S200E had greater intrinsic FRET efficiency with the fluorophores moving ∼3 Å closer together in HEK293T cells (Fig. 2H). Dimer formation was also decreased demonstrating that ERβ–S200E had a lower affinity for ERβ interactions compared to the phosphonull (Fig. 2I, J). There were no observed structural changes for the ERβ–S200 phosphomutants in SK–N–SH cells resulting in equivalent intrinsic FRET efficiency between the ERβ–S200A and S200E (Fig. 2K). However, there were significantly fewer ERβ–S200E dimers compared to ERβ–S200A in SK–N–SH cells, demonstrating decreased affinity (Fig. 2L, M). It is also important to note that overall ERβ dimer formation was much higher for both phosphonull and phosphomimetic mutants in the SK–N–SH neuronal cells (Fig. 2D, G) compared to HEK293T cells (Fig. 2J, M), which was consistent with our results for ERβ–WT in that cell type. Taken together, these results indicate that phosphorylation at both sites in the ERβ DBD would induce structural changes and decrease the affinity of ERβ–ERβ interactions in HEK293T cells. Phosphorylation in the ERβ DBD in neuronal cells would also decrease ERβ–ERβ affinity, but the decreased affinity would be unlikely due to changes in structural conformation.

### Phosphorylation of ERβ–Y488 on helix 12 decreased ERβ:ERβ dimerization (Fig. 3)

Our previous study using AQUA–MS also quantified ERβ phosphosites that were located outside of the DBD. Of particular interest was tyrosine 488 (Y488) located at the at the far N–terminal side of H12 in the ligand binding domain (LBD) (Fig. 3A). This amino acid is equivalent to ERα–Y537, which is often mutated in metastatic breast cancer [18-21]. Cell–free biochemical assays have demonstrated that ERβ–Y488 phosphorylation can alter co–activator protein recruitment to the AF–2 groove and potentially facilitate ERβ ligand–independent activity [22]. Our *in vivo* study showed that ERβ–Y488 was highly phosphorylated in the human brain (∼60%), not age–dependent, and it did not impact ERβ DNA binding ability or transcriptional activity [14]. However, the functional consequences of ERβ–Y488 phosphorylation have not been tested in any cell type.

**Figure 3.**
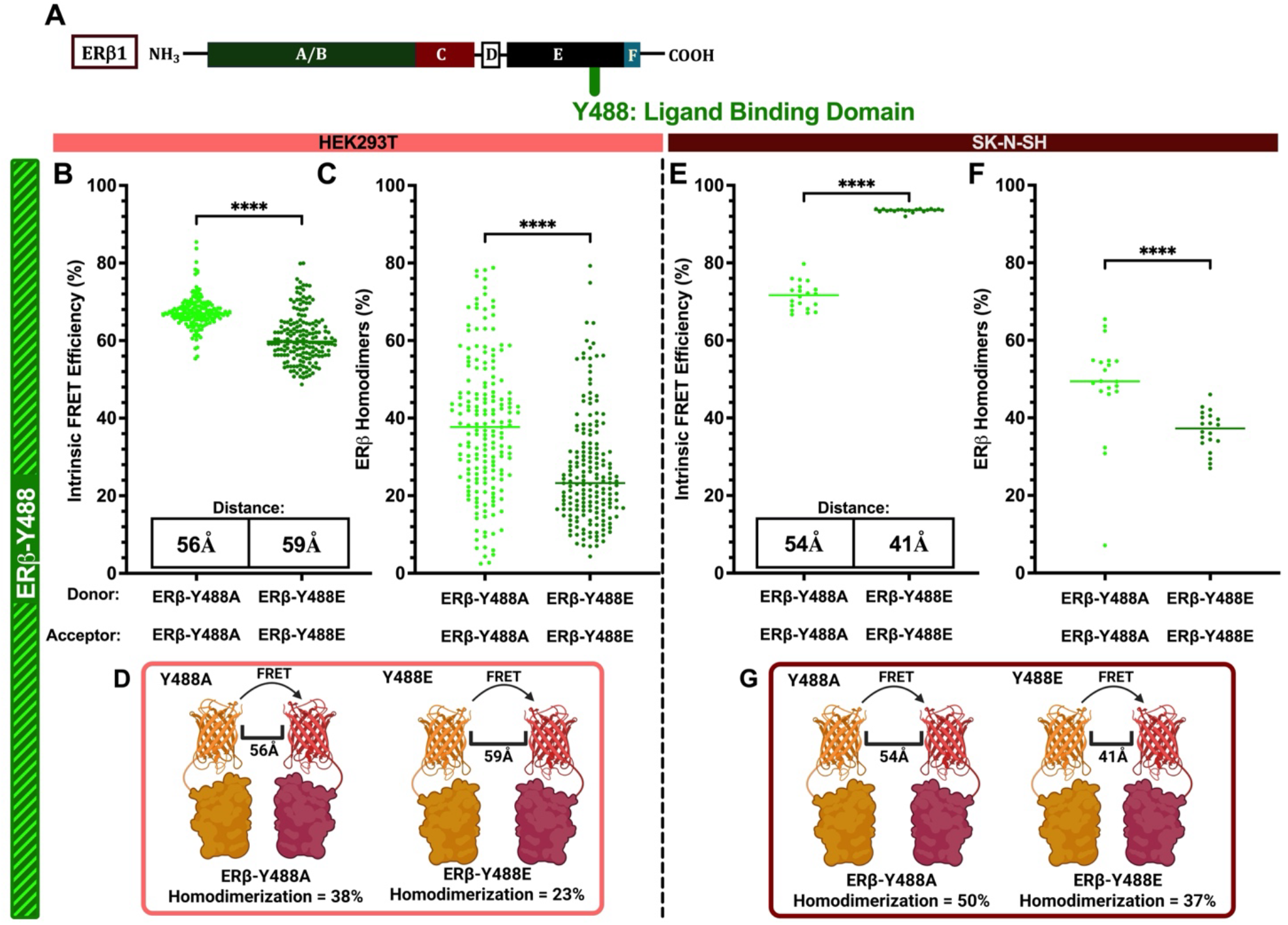
Phosphorylation of ERβ-Y488 decreased ERβ:ERβ dimerization. (A) Diagram depicting ERβ protein domains and the relative location of Y488 that was mutated to phosphonull (S>A) or phosphomimetic (S>E). Intrinsic FRET efficiency, fluorophore distance, and homodimer formation comparing Y488A to Y488E (green) were calculated using time-correlated single-photon counting (TCSPC) in HEK293T (B, C) and SK-N-SH cells (E, F). Cartoon depictions of quantified data are illustrated in panels D, G. Data are displayed as median ± SD. Data were collected across 4 different biological replicates and 2 independent transfections per biological replicate. A non-parametric Kolmogorov Smirnov test was used for statistical analysis. An **** denotes p<0.0001. Created in BioRender. Yildiz, Y. (2026) https://BioRender.com/e6c5www

Our results showed that intrinsic FRET efficiency was decreased for Y488E compared to Y488A, with a corresponding structural shift of increased fluorophore distance of ∼3 Å in HEK293T cells (Fig. 3B). ERβ–Y488E also formed fewer dimers, which demonstrated decreased affinity compared to ERβ–Y488A (Fig. 3C, D). Notably, we observed a very large structural change for ERβ–Y488E in SK–N–SH cells resulting in a 13 Å decrease in fluorophore distance compared to ERβ–Y488A, which corresponded to a very high intrinsic FRET efficiency of 93% (Fig. 3E). These results raise the possibility that phosphorylation of ERβ–Y488 would induce conformational changes that are large enough to impact the accessibility of ligands and/or coregulatory proteins for binding ERβ in neuronal cells. Moreover, ERβ–Y488E formed significantly fewer dimers in SK–N–SH cells, similar to the other phosphomimetics in both cell types, suggesting that phosphorylation decreases affinity and that this is a conserved feature across cell types (Fig. 3D, G).

### E2 increased ERβ dimer formation regardless of phosphorylation state (Figs. 4, 5)

Next, we tested the effects of E2–mediated ERβ dimerization for each phosphomutant in the ERβ DBD. Briefly, HEK293T cells were grown in charcoal–stripped FBS (i.e. apo ERβ) or in charcoal–stripped FBS supplemented with 10 nM E2. HEK293T cells were chosen for these experiments to 1) minimize the confounding endogenous factors mediating E2 signaling that are present in SK–N–SH cells, and 2) compare directly with our results from ERβ–WT after E2 treatment in HEK293T cells. As we showed in those experiments, E2 treatment increased ERβ-WT dimerization and induced an expected conformational change (see Fig. 1). Similarly, experiments testing ERβ phosphomutants revealed E2 treatment increased dimer formation for all phosphomutants demonstrating that ligand binding can induce dimer formation regardless of the phosphorylation state of these amino acid residues (Figs. 4, 5).

**Figure 4.**
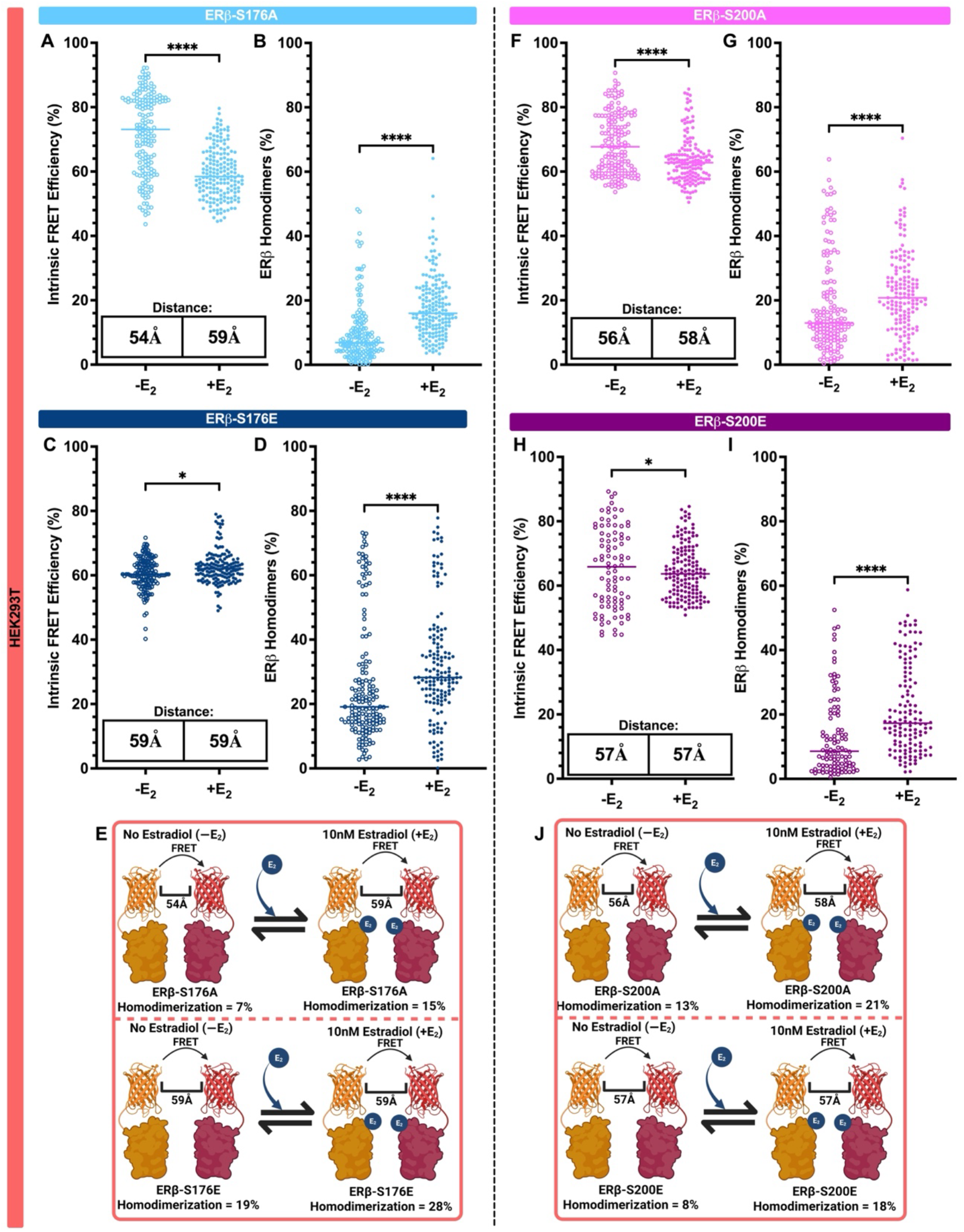
E2 increased ERβ dimer formation regardless of phosphorylation state of the DNA binding domain. Intrinsic FRET efficiency, fluorophore distance, and homodimer formation comparing S176A (A and B) and S176E (C and D) following 10 nM E2 treatment were calculated using time-correlated single-photon counting (TCSPC) in HEK293T cells. Panels F-I show the same parameters for S200 A (F and G) and S200E (H and I). Cartoon depictions of quantified data are illustrated in panels E and J. Data are displayed as median ± SD. Data were collected across 4 different biological replicates and 2 independent transfections per biological replicate. A non-parametric Kolmogorov Smirnov test was used for statistical analysis. An * denotes p<0.05, **** p<0.0001. Created in BioRender. Yildiz, Y. (2026) https://BioRender.com/lm85jzc

Specifically, E2 treatment decreased intrinsic FRET efficiency of ERβ–S176A due to a significant increase in fluorophore distance of 5 Å, and also increased dimer formation (Fig. 4A, B, E). On the other hand, there was minimal effect of E2 on fluorophore distance for ERβ–S176E, yet we still observed a statistically significant increase in intrinsic FRET efficiency (Fig. 4C). Moreover, E2 increased dimer formation of ERβ–176E similar to what we observed with phosphonull (Fig. 4D, E). These results were very similar for ERβ–S200 in that both phosphomutants had minimal structural shifts, small changes in FRET efficiency, and formed more dimers in response to E2 treatment (Fig. 4F–J), suggesting that phosphorylation at this site is not an important driver of E2-mediated ERβ dimerization.

Also of note, ERβ–S176A in the apo state had very low levels of dimerization (7%) and E2 treatment only increased it to 15% (Fig. 4B). This was contrasted with significantly higher dimerization in the apo state for the phosphomimetic at that same site (ERβ–S176E = 19%), which then increased to 28% following E2 treatment (Fig. 4D). These data show that the magnitude of E2–induced dimer formation can depend on the phosphorylation state of ERβ–S176 in the DBD. Our previous study showed that phosphorylation of ERβ–S176 decreased with age and these data demonstrate that an age-dependent loss of ERβ–S176 phosphorylation could have important biological consequences in the context of hormone replacement therapy.

We hypothesized that E2 would have the greatest impact on structural changes and dimerization when ERβ–Y488 was phosphorylated due to its proximity to helix 12. Contrary to that expectation, E2 did not induce large structural changes when ERβ–Y488 was mutated to a phosphonull or phosphomimetic (Fig. 5A, C). Molecular dynamic modeling predicted that H12 would be in an agonistic conformation when ERβ–Y488 was phosphorylated [22], and our results suggest that ligand binding would not further alter that conformation. However, E2 increased dimer formation for both ERβ–Y488A (Fig. 5B) and ERβ–Y488E (Fig. 5D), suggesting that phosphorylation at this site is not important for E2–mediated ERβ–ERβ affinity (Fig. 5E). Notably, the amount of dimer formation in the apo state was very low for most of the phosphomutants, which was similarly observed with ERβ–WT. An exception to this was ERβ–Y488A, which had the highest percentage of dimers formed in the apo state (29%, Fig. 5B) compared to all other phosphomutants and was nearly equivalent to ERβ–WT in normal growth media in HEK293T cells.

**Figure 5.**
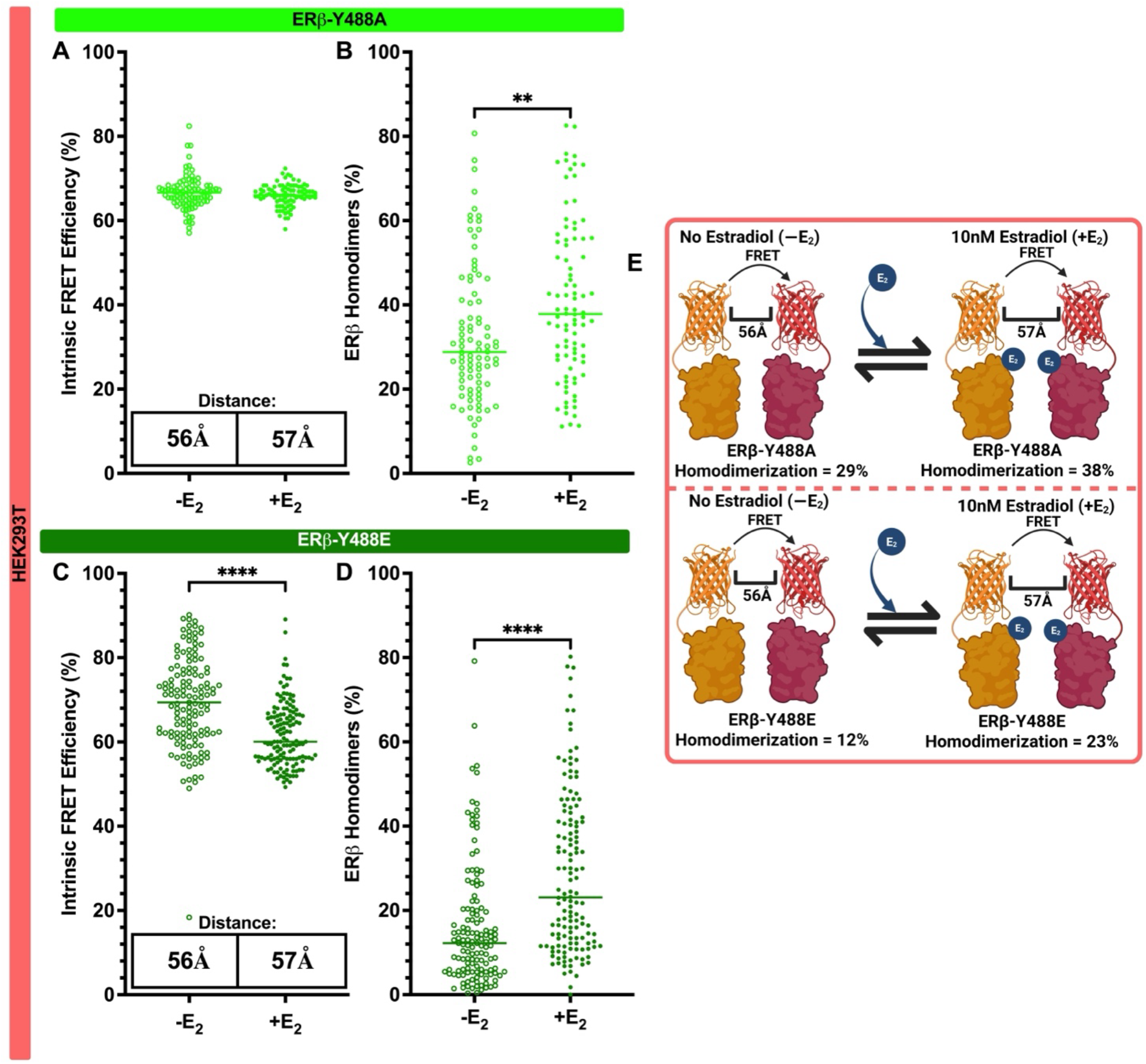
E2 increased ERβ dimer formation regardless of phosphorylation state of the ligand binding domain. Intrinsic FRET efficiency, fluorophore distance, and homodimer formation comparing Y488A (A and B) and Y488E (C and D) following 10 nM E2 treatment were calculated using time-correlated single-photon counting (TCSPC) in HEK293T cells. vCartoon depictions of quantified data is illustrated in panel E. Data are displayed as median ± SD. Data were collected across 4 different biological replicates and 2 independent transfections per biological replicate. A non-parametric Kolmogorov Smirnov test was used for statistical analysis. An ** denotes p<0.01, **** p<0.0001. Created in BioRender. Yildiz, Y. (2026) https://BioRender.com/0qg8bc7

### ERβ phosphorylation impairs ERβ–mediated repression of the amyloid precursor protein (APP) promoter (Fig. 6)

Our data thus far has shown that phosphorylation of ERβ–S176, –S200, and – Y488 can regulate ERβ homodimer formation in a cell type specific manner. However, the consequences of these biophysical interactions to downstream ERβ–mediated signaling and transcriptional regulation remain unknown. Previously, we showed that ERβ repressed human amyloid precursor protein (APP) promoter activity [6]. Based on these results, we hypothesized that differential protein interactions between the phosphonull and phosphomimetic ERβ mutants would alter ERβ–mediated transcriptional regulation of the APP promoter. To test this hypothesis, we co–transfected neuronal SK–N–SH cells with the human APP–luciferase promoter along with our phosphonull and phosphomimetic ERβ mutants and quantified promoter activity using luciferase assays, as in our previous study [6]. Consistent with our prior study using ERβ–WT, all ERβ phosphomutants repressed APP promoter activity to some degree. Specifically, ERβ–S176A and S176E repressed APP promoter activity equivalently (45%, 53% respectively, Fig. 6A). By contrast, ERβ–S200A decreased APP promoter activity by 55%, compared to only 25% repression by S200E (Fig. 6B). ERβ–Y488 was similar to ERβ–S200 with a 62% decrease in promoter activity for the phosphonull, Y488A, compared to only 35% for the phosphomimetic Y488E (Fig. 6C). Taken together, these results corroborate our FRET results showing that site specific phosphorylation states in the ERβ DBD and H12 impact ERβ–protein interactions and their ability to regulate transcription at an ERβ target gene promoter.

**Figure 6.**
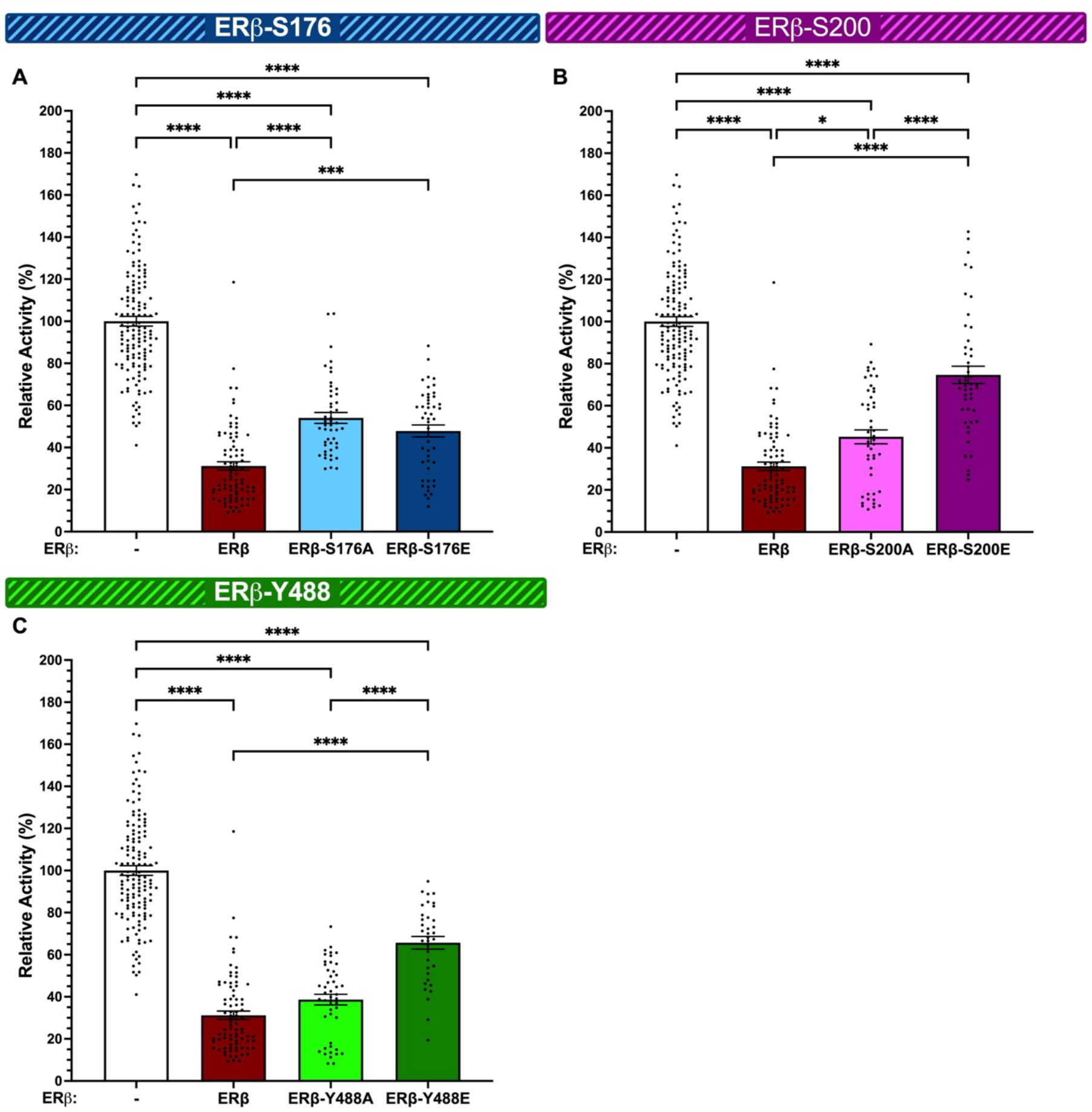
ERβ phosphorylation of S200 and Y488 impairs ERβ-mediated repression of the amyloid precursor protein (APP) promoter. Human amyloid precursor protein (APP) promoter activity was measured by luciferase assays in SK-N-SH cells following transient transfection with a reporter plasmid vector containing APP-luc and expression plasmid vector containing ERβ phosphomutants at S176 (A), S200 (B) and Y488 (C). Relative promoter activity was calculated based on % change from APP-luc promoter alone. Data were averaged from at least 3 independent biological replicates with 12 technical replicates per biological replicate. Data are depicted as mean ± SEM. One-way ANOVA was used for statistical analysis. An * denotes p<0.05, *** p<0.001, **** and p< 0.0001.

## DISCUSSION

The importance of age–dependent posttranslational modifications to ERβ are poorly described, and our data support the premise that these posttranslational modifications likely contribute to major shifts in ERβ signaling across the lifespan. Phosphorylation of amino acids in the N–terminal A/B domain are the most well–studied posttranslational modifications on ERβ. The A/B domain encompasses amino acids 1–148 and harbors an activator function–1 (AF–1) region that can induce ligand–independent activity of ERβ [23-28]. Peptide growth factors, such as epidermal growth factor (EGF), trigger kinases that phosphorylate S87 and S105 in the AF–1 region, thereby enabling the recruitment of co–activator proteins leading to ERβ–mediated activation of target genes [12, 29, 30]. Here, we provide evidence that phosphorylation of amino acids outside of the well–characterized ERβ AF–1 region caused structural shifts that altered the ability of ERβ to form homodimers. Specifically, we demonstrated that 1) phosphorylation of ERβ–S176, –S200, and –Y488 reduced ERβ homodimer formation regardless of cell type, 2) ERβ homodimer formation was strengthened in a neuronal cell type compared to HEK293T cells, and 3) ERβ phosphorylation at ERβ–S200 and ERβ–Y488 resulted in a partial loss of ERβ–mediated APP promoter gene repression.

In this study we interrogated two previously uncharacterized ERβ amino acids in the C domain of ERβ. This domain, known as the DNA–binding domain (DBD), extends from amino acids 149 to 214 and consists of two parallel zinc fingers (aa 149–169 and 185–209, respectively) that together facilitate ERβ–DNA binding [31-34]. S176 lies just distal to the first zinc finger, which recognizes specific base pair sequences on ERβ target genes. Our previous work showed that phosphorylation of S176 decreased with age in the human brain and impaired DNA binding [14]. ERβ binds DNA as a dimer and the data herein demonstrated that S176 phosphorylation decreased dimer formation, which would predict a reduced ability to bind DNA consistent with our prior study. On the other hand, S200 lies within the second zinc finger of the DBD. The second zinc finger stabilizes ERβ–DNA interaction through nonspecific binding to the DNA backbone. Using AQUA–MS we quantified phosphorylation of S200 and showed that it had the lowest level of phosphorylation compared to the other sites tested and, similar to S176, there was a significant age-dependent decrease in phosphorylated S200 coupled with a complete abolishment of DNA binding ability [14]. The data shown herein demonstrate that the ERβ phosphomimetic S200E reduced ERβ dimer formation providing a mechanistic explanation for its inability to bind DNA. Furthermore, our luciferase data showed decreased ability of ERβ–S200E to repress the APP promoter compared to its phosphonull counterpart, consistent with our previous work and current findings of reduced ERβ dimer formation of ERβ–S200E. Although the results for S176 and S200 provide a mechanistic basis for our previous findings, it was unexpected that phosphorylation within the DBD would impair ERβ dimerization. The DBD is predicted to be a weak dimerization interface for estrogen receptors with the primary dimer interface dependent on H10/H11 contacts in the LBD, which are strengthened upon ligand binding [35]. The DBD forms intermolecular contacts through a well conserved 5 aa region in the second zinc finger known as the D box. These contacts are formed once the receptor binds to response elements on specific DNA targets. The presence of a D box for ERβ has been assumed based on homology with other nuclear receptors but has not been experimentally confirmed to date. Our data are consistent with the presence of a functional D box in the ERβ DBD that is required for dimer formation, and tunable by age-dependent posttranslational modifications.

There is high homology between the DBD among members of the nuclear receptor family making it likely that our data are broadly applicable to how aging and/or phosphorylation impacts other nuclear steroid hormone receptors. Indeed, the equivalent site for ERβ–S200 on ERα is S236 and phosphomimetic mutation (S236E) reduced ERα homodimer formation [36]. Interestingly, phosphomimetic mutation also abolished heterodimer formation of ERα with ERβ, but the phosphorylation state of ERβ in that experiment was not reported. Similarly, phosphorylation of the farsenoid X receptor (FXR) DBD at S135 and S154 increased coactivator recruitment and ligand–dependent activation of the receptor. FXR–S154 is located between the two zinc fingers, which somewhat corresponds to the location of ERβ-S176 [37-40]. FXR forms heterodimers with the retinoic acid receptor (RXR), but that interaction was not affected by FXR phosphorylation of the DBD. The dimerization of many nuclear receptors is stabilized by residues in the hinge region (i.e. D domain) and previous studies have demonstrated that phosphorylation of mouse ERβ–S255 (equivalent to human D236) in the hinge region reduced E2–mediated activity of ERβ, but whether the effect was due to impaired homodimer formation was not tested [41].

We also investigated how phosphorylation of ERβ–Y488 altered ERβ–protein interactions. This amino acid is located at the far end of the C–terminus in the ligand–binding domain (LBD = aa 294-498). The LBD is critical for ERβ homo– and hetero–dimerization whereby helix 11 (H11) from each monomer interact through conserved hydrophobic residues at the N–terminus, supported by residues of H8 and the loop between H9 and H10 [17, 42-44]. Upon E2 binding, H12 rotates perpendicular to the dimerization area and forms a “lid” on the ligand binding cavity. This ligand–induced structural change exposes the ligand–dependent activation function 2 (AF–2) surface, which allows recruitment of specific coactivators at the LXXLL interface and functionally transitions ERβ to a "transcriptional activator" at select target gene promoters [45, 46]. Y488 is positioned at the junction between H11 and H12 outside of the predicted intermolecular surfaces required for dimer formation. Our prior results showed that phosphorylation of Y488 was remarkably stable across the lifespan and was not affected by E2 binding to the LBD [14]. Further, our AQUA–MS data showed that approximately 60% of ERβ–Y488 residues were phosphorylated in women’s brains at all ages tested and changing the phosphostate did not alter ERβ–DNA binding, so the functional significance of this phosphorylation site was unclear. In this study, we observed that the phosphomimetic ERβ–Y488E decreased ERβ–ERβ dimer formation, similar to ERβ–S176E and ERβ–S200E, and the effect was independent of cell type. Moreover, ERβ–Y488E was less effective at repressing APP promoter activity compared to ERβ–Y488A. Together, these data suggest the possibility that ERβ–Y488 functions as a tunable molecular rheostat, and the approximately 60% observed basal phosphorylation at this site establishes a dynamic equilibrium between receptor dimer formation and transcriptional activity. This idea is consistent with Tharun and colleagues who used molecular dynamic modelling to demonstrate that phosphorylation of apo–ERβ Y488 could induce unfolding of H12 causing it to partially fold back on the ligand binding domain [22, 47]. This conformational change moves H12 to the "agonist" position independent of ligand binding, thereby exposing the LXXLL coactivator docking motif/AF–2 surface and shifting the receptor towards transcriptional activation instead of repression.

Phosphomimetic and phosphonull mutants are useful tools to study the functional consequences of site–specific phosphorylation of proteins, however it is important to note that amino acid substitutions only partially mimic phosphorylation properties. Phosphates introduce a bulky negatively charged polar group to amino acids, which can alter overall shape and intramolecular interactions. Glutamic acid substitution of a phosphoserine results in a less negatively charged residue, which might not fully replicate functional changes that would occur with the native phosphoserine. Similarly, glutamic acid substitution of a phosphotyrosine can create a similarly electronegative condition but the location of the modification is not equivalent to adding a phosphate group. Additionally, post translation modifications do not occur in isolation at single sites. ERβ can be targeted by multiple kinases determined by cell type and intracellular milieu. The amino acids are phosphorylated in concert, and thus single phosphorylated sites likely do not depict the global phosphorylated state of ERβ. Despite these limitations, understanding the contribution and relative impact of single site PTMs can uncover mechanistic insight on how cell-specific factors and cellular microenvironments alter ERβ structure, ERβ–protein interactions and downstream signaling.

Overall, our data revealed that phosphorylation of previously uncharacterized amino acids in the DBD decrease ERβ–ERβ dimer formation and functional activity at an ERβ target gene promoter. Future studies are aimed at elucidating the kinases and phosphatases that contribute to the age-dependent phosphostate of ERβ, which could uncover new therapeutic strategies for neurological diseases and cognitive deficits associated with menopause.

## METHODS

### Cell Culture

SK-N-SH cells, a human neuroblastoma cell line isolated from a four-year-old female were purchased from American Type Culture Collection (ATCC, Manassas, VA, USA) and maintained in the recommended normal growth media of minimal essential media with Earle’s balanced salts (MEM/EBSS-Cytiva, MA, USA. SH30244.01) plus 1% L-glutamine, 1% non-essential amino acids (Sigma-Aldrich, M7145) and 10% heat-inactivated fetal bovine serum (FBS) (BenchMark™ GeminiBio, Cat# Cat#100-106) and Plasmocin™ prophylactic (25 mg/500 ml; Invivogen, ant-mpp). HEK293T cells, a derivative of the human embryonic kidney cell line (HEK293), was purchased from American Type Culture Collection (ATCC, Manassas, VA, USA) and maintained in the recommended normal growth media of Dulbecco’s modified Eagle medium (DMEM), 10% heat-inactivated fetal bovine serum (FBS) (BenchMark™ GeminiBio, 100-106) and Plasmocin™ prophylactic (25 mg/500 ml; Invivogen, ant-mpp). Cells were grown to 60-70% confluency prior to all experiments. All experiments were performed within 10 passages, and different passages were considered to represent biological replicates. HEK293T and SK-N-SH cells do not express endogenous ERβ, as validated previously.

### Cell Treatment

Normal growth media (described above) was replaced with media containing phenol red-free MEM or DMEM, the appropriate supplements and 10% charcoal/dextran stripped FBS (GeminiBio, Sacramento, CA, USA. Cat #100-106-500) for at least 24 hours to eliminate all sources of hormones within the media. Cells were treated with 10 nM 17β-estradiol (E2; Steraloids) or an equivalent volume of media containing 0.001% ethanol (vehicle control) overnight for 16 hours.

### Plasmid Constructs

The plasmid expression vector containing an insert for human ERβ1 was generously provided by Dr. Shuk-Mei Ho (University of Cincinnati, Cincinnati, OH, USA) and has been extensively characterized [48]. mCyRFP1 tag and mMaroon1 empty plasmid vectors were obtained from Addgene and have previously been characterized for use in FLIM–FRET [49-52]. The full ERβ coding sequence was cloned into the mCyRFP1 or mMaroon1 tag vector using Gibson assembly resulting in N- and C-terminal-fused ERβ with the respective tags. Subsequently, site-directed mutagenesis was used to create phosphomutants for S176, S200, and Y488 using the ERβ-tagged mCyRFP1 or mMaroon1 tagged ERβ (Q5 site–directed mutagenesis kit, E0554S, New England Biolabs, Inc., MA, USA). Primers were designed using the NEBase changer primer design tool available from New England Biolabs website. Single amino acid changes (i.e. serine/tyrosine > glutamic acid or alanine) were created following manufacturer instructions and amino acid switch was verified by DNA sequencing (ACGT Inc, IL, USA). The human APP promoter report vector (GoClone, #S719381) was purchased from SwitchGear Genomics (Menlo Park, CA, USA). Sequences for all plasmids were verified by DNA sequencing (ACGT, Inc., Wheeling, IL, USA).

### Transient transfections

Cells were seeded at 2.0 x 10^5^ cells per well in a 6 well plate 24 hours prior to transfection. All plasmids were transfected in replicate wells of 2 minimum/construct/condition and experiments were repeated a minimum of 4 times using separate passages, but within 10 passages of each other. SK-N-SH cells were transfected using the lipid-mediated transfection reagent Fugene 4K (Fugene LLC, WI, USA, cat. #4K-1000). Expression vectors were transfected at a ratio of 1000 ng:6.0 µl (DNA:Fugene). HEK293T cells were transfected using the lipid-mediated transfection reagent Lipofectamine 3000 (Invitrogen, cat #L3000075). Expression vectors were transfected at a ratio of 60 ng:1.5 µl:1.0 µl (DNA:P3000:Lipofectamine). mCyRFP1 (donor) and mMaroon (acceptor) constructs were co-transfected at a 1:5 ratio.

### Fluorescence Lifetime Measurements

Fluorescence lifetime measurements were performed using time-correlated single-photon counting (TCSPC) as previously described [53-55]. TCSPC histograms were acquired for cells expressing ERβ-WT-mCyRFP1, S176A, S176E, S200A, S200E, Y488A or Y488E mutants. Cells expressing both mCyRFP1 and mMaroon1 constructs were manually selected using a defocused excitation supercontinuum laser with 20 MHz repetition frequency (SuperEvo K, NKT Photonics, Denmark) to excite the whole cell. Donor (mCyRFP1) signal was acquired using an excitation filter 482/12-nm bandpass filter with 1.0 neutral density filter. Fluorescence emission was detected using a 640/50-nm emission bandpass filter. After cell selection using the defocused laser beam, the defocusing lens was removed from the light path, the laser intensity was attenuated with a 0.5 neutral density filter, and the focused laser was placed on the cell yielding 100,000 (HEK293T cells) – 50,000 (SK-N-SH cells) photons/s count rate. Fluorescence was detected through 1.2 NA water immersion objective with an PMA hybrid detector connected to a photon-counting module (HydraHarp 300, PicoQuant, Germany) with a time channel width of 16 ps.

HEK293T cells expressing mCyRFP1-labelled ERβ and mMaroon1-labeled ERβ showed multiexponential decays with two components. To quantify these components, fluorescence decay histograms from four independent transfections per biological replicate were analyzed in SymPhoTime 64 software using global analysis. The intensity decays were fitted with a multiexponential model using the formula:

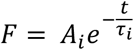

where i = 1,2,…,n. Here, F is fluorescence intensity, A_i_ are amplitudes of i components, and t_i_ are lifetimes of each component. Specifically, for HEK293T cells A_1_ and A_2_ are amplitudes of two components and τ_1_ and τ_2_ are fluorescent lifetimes of donor alone and lifetime of FRET species.

### Luciferase Assay

Luciferase assays were conducted as previously described [6]. Briefly, SK-N-SH cells were seeded at 5,000 cells per well in 96-well plates with white bottoms. 24 hours after seeding, cells were co-transfected with the APP-luc reporter and ERβ WT or phosphomutant expression vectors. 48 hours after transfection, plates were stored in −80°C overnight. Plates were thawed on wet ice before the addition of LightSwitch Assay Reagent per manufacturer instructions (LightSwitch, cat #LS100), and luciferase activity was measured using a CLARIOstar microplate reader (BMG Labtech, NC, USA).

## Notes

### Competing Interest Statement

The authors have declared no competing interest.

### Funding

- National Institutes of Health: 5R01AG082135

